# MntJULiP and Jutils: Differential splicing analysis of RNA-seq data with covariates

**DOI:** 10.1101/2024.01.01.573825

**Authors:** Wui Wang Lui, Guangyu Yang, Liliana Florea

**Affiliations:** Department of Computer Science, Johns Hopkins University, Baltimore MD 21205; TikTok, 1199 Coleman Ave, San Jose, CA 95110; Department of Genetic Medicine, Johns Hopkins School of Medicine, Baltimore, MD 21205

## Abstract

Differences in alternative splicing patterns can reveal important markers of phenotypic differentiation, including biomarkers of disease. Emerging large and complex RNA-seq datasets from disease and population studies include multiple confounders such as sex, age, ethnicity and clinical attributes, which demand highly specialized data analysis tools. However, few methods are equipped to handle the new challenges. We describe an implementation of our programs MntJULiP and Jutils for differential splicing detection and visualization from RNA-seq data that takes into account covariates. MntJULiP detects intron-level differences in alternative splicing from RNA-seq data using a Bayesian mixture model. Jutils visualizes alternative splicing variation with heatmaps, PCA and sashimi plots, and Venn diagrams. Our tools are scalable and can process thousands of samples within hours. We applied our methods to the collection of GTEx brain RNA-seq samples to deconvolute the effects of sex and age at death on the splicing patterns. In particular, clustering of covariate adjusted data identifies a subgroup of individuals undergoing a distinct splicing program during aging. MntJULiP and Jutils are implemented in Python and are available from https://github.com/splicebox/.

## Introduction

Differences in alternative splicing patterns are responsible for the diversity of proteins across tissues, cell types and developmental stages, and disruptions in normal RNA splicing patterns have been reported in a number of diseases. Increasingly large and complex RNA-seq datasets that include multiple confounders such as sex, age, ethnicity and clinical attributes, are emerging from disease study cohorts and population-level projects, which demand highly specialized analysis tools. While multiple methods exist to detect differences in splicing, including LeafCutter, MntJULiP, rMATS, SUPPA2 and DARTS [1-5], few are equipped to handle the complexities of the data. In particular, there is a scarcity of programs that can rigorously account for the effect of confounding attributes on the observed data. Additionally, visualization tools are critical in enabling biologists to quickly and intuitively interpret such differences, to identify patterns genome-wide or at the level of the individual genes. We previously developed two tools, MntJULiP [2] and Jutils [6], for differential splicing detection and visualization, respectively, that can efficiently handle large scale and complex RNA-seq data collections. We report here a recent implementation of these programs to account for covariates and to produce new customizable visualizations of covariate adjusted data as PCA plots. We demonstrate their accuracy on simulated data. We then illustrate their applicability and usefulness with an analysis of RNA-seq data from brain tissue obtained from the GTEx repository [7], which paints the global splicing landscape across age and biological sex groups and identifies a potential distinct splicing program in a subset of individuals.

### Differential splicing (DS) detection with MntJULiP

MntJULiP [2] detects differences in splicing at the intron level, which greatly improves performance when transcript reconstructions are inaccurate or incomplete. It identifies events directly from the alignment data, without relying on a reference gene annotation, and therefore can find and report novel events. MntJULiP can detect both differences in the introns’ splicing ratios (DSR), and changes in the abundance level of introns (DSA), and thus can capture alternative splicing variations in a comprehensive way. For DSA, it considers each intron individually and models read counts with a zero inflated negative binomial (ZINB) distribution. For DSR, it groups introns sharing a splice site into a ‘bunch’, and uses a Dirichlet multinomial (DM) distribution to simultaneously model all introns in a group (**Supplementary Fig. S1**). Additionally, MntJULiP has the rare ability to perform multiple comparisons simultaneously, which we showed was more accurate at capturing global differences in a time series or complex experiments [2].

### Differential splicing visualization with Jutils

Jutils [6] is a Python package for visualization of differential splicing events in the form of heatmaps, sashimi plots and Venn diagrams of sets of differentially spliced genes. Jutils can be used with virtually any differential splicing detection tool, and is specifically configured to work with the output of popular methods including LeafCutter, MntJULiP and rMATS. It converts the output of each tool into an intermediate TSV formatted file to render the data in a unified format, described in [6]. This file is then sufficient to create visualizations, making it lightweight and very well suited for collaborations.

We recently introduced visualizations of PCA plots to identify potential unknown relationships among samples, and to observe the effect of covariates on the data. PCA plots are generated from the splicing ratios (PSI = ‘Percent Splice In’) for DSR methods, or read counts for DSA methods, which are contained in the TSV file and imported from the output of differential splicing programs. As customizable features, input data can be filtered by significance, and data points can be differentiated by color, shape and label based on conditions or covariates.

### Covariate augmented Bayesian models

We introduce covariate effects as linear components in the MntJULiP Bayesian models to account for extraneous attributes that could bias the analysis (described in the **Supplementary Method M1)**. Numerical covariates, such as age and weight, are centered and scaled, while categorical covariates, such as biological sex and ethnicity, are encoded into a numerical representation (0, 1, … K-1). Using the covariate augmented models, we generate new adjusted counts and PSI values per sample, which can be used to generate global views of the alternative splicing variation as heatmaps or PCA plots using Jutils or other visualization tools.

To validate the covariate models, we simulated data for one ‘condition’, with values ‘control’, ‘disease’ and ‘stage2’, with one covariate, ‘biological sex’, with values ‘M’ and ‘F’. For each condition, we generated 5 samples in each of the categories ‘condition’*x*’biological sex’. Starting from an empirical transcript expression matrix trained on an RNA-seq data set from lung fibroblasts (GenBank A# SRR493366) and using GENCODE v.41 as reference, we generated 11.5 million 100 bp long paired-end reads per sample from 2,000 genes with two or more expressed isoforms. Changes in splicing were simulated as described in [2], separately for the DSR and the DSA models; detailed descriptions are provided in the **Supplementary Method M2**.

We evaluated the performance of MntJULiP in pairwise splicing ratio (DSR) comparisons, with and without covariates, alongside LeafCutter, the only other program to implement covariates, on the simulated data in an imbalanced comparison of (4M,1F) ‘control’ versus (4F,1M) ‘disease’ samples (**Supplementary Figure S2**). MntJULiP retains the performance of the original implementation and drastically reduces the false positives due to covariate bias. Also, MntJULiP has consistently better performance over LeafCutter, as measured by the F-value. Further, the PCA plots of the estimated PSI values indicate that variation due to ‘biological sex’ was correctly removed. Similar results were observed for pairwise splicing abundance (DSA) comparisons (**Supplementary Figure S3**), and for the DSR and the DSA multiway comparisons (**Supplementary Figure S4**). Therefore, MntJULiP accurately models and removes biases due to covariates form the RNA-seq data. Notably, MntJULiP is the *only* tool with this capability for differential splicing abundance (DSA) applications, and also the *only* tool available for multiway comparisons with covariates.

### Deconvoluting the effects of covariates on human frontal cortex splicing from GTEx RNA-seq data

To illustrate, we applied our methods to 1,398 GTEx RNA-seq samples from 13 brain regions, in three comparisons. The first comparison, among regions, revealed distinct groupings between the cerebellar, cortex, and basal ganglia regions, which did not change when accounting for the covariates ‘biological sex’ and ‘age at death’ (**Supplementary Figure S5**). Secondly, we compared 120 frontal cortex RNA-seq samples by age groups (‘20s’, …, ‘70s’). Changes in frontal cortex in aging are thought to contribute to sex-specific differences in the prevalence of neurological disorders [8]. More differences were observed with more distant age groups, consistent with reports of splicing deregulation with aging [9] (**Figure 1A**). As an outlier, the larger number of events between the ‘20s’ and ‘30s’ groups may be due to the small number of samples in those categories (3 and 4, respectively). When ‘biological sex’ was used as a covariate, a significant increase in the number of differences was observed between the ‘20s’ and ‘40s’ groups, indicating a possible mark of sex-specific differentiation. Lastly, we compared the 83 male (‘M’) and 37 female (‘F’) frontal cortex samples to identify sex-specific differences in splicing. Jutils heatmaps of PSI values revealed a subgroup of 22 samples (12 ‘F’ and 10 ‘M’) with a distinct alternative splicing pattern, which became evident when regressing for ‘age at death’. This subgroup was higher aged (between 50-79 years of age), had an over-representation of females (12F:10M compared to 37F:83M for the entire set), and pointed to a probable distinct splicing program in a subset of individuals with aging (**Figure 1,B and C**). Of note, similar DSA analyses of the frontal cortex data show a clear separation of samples by ‘biological sex’, with or without covariate adjustment, and reveal distinguishing events including at the non-coding X Inactive Specific Transcript (XIST) gene involved in X chromosome inactivation in female early development processes (**Supplementary Figure S6**). Therefore, as we previously noted [2], DSR and DSA reflect different and complementary views and effects of alternative splicing on the transcriptional and functional outcomes.

**Figure 1.**
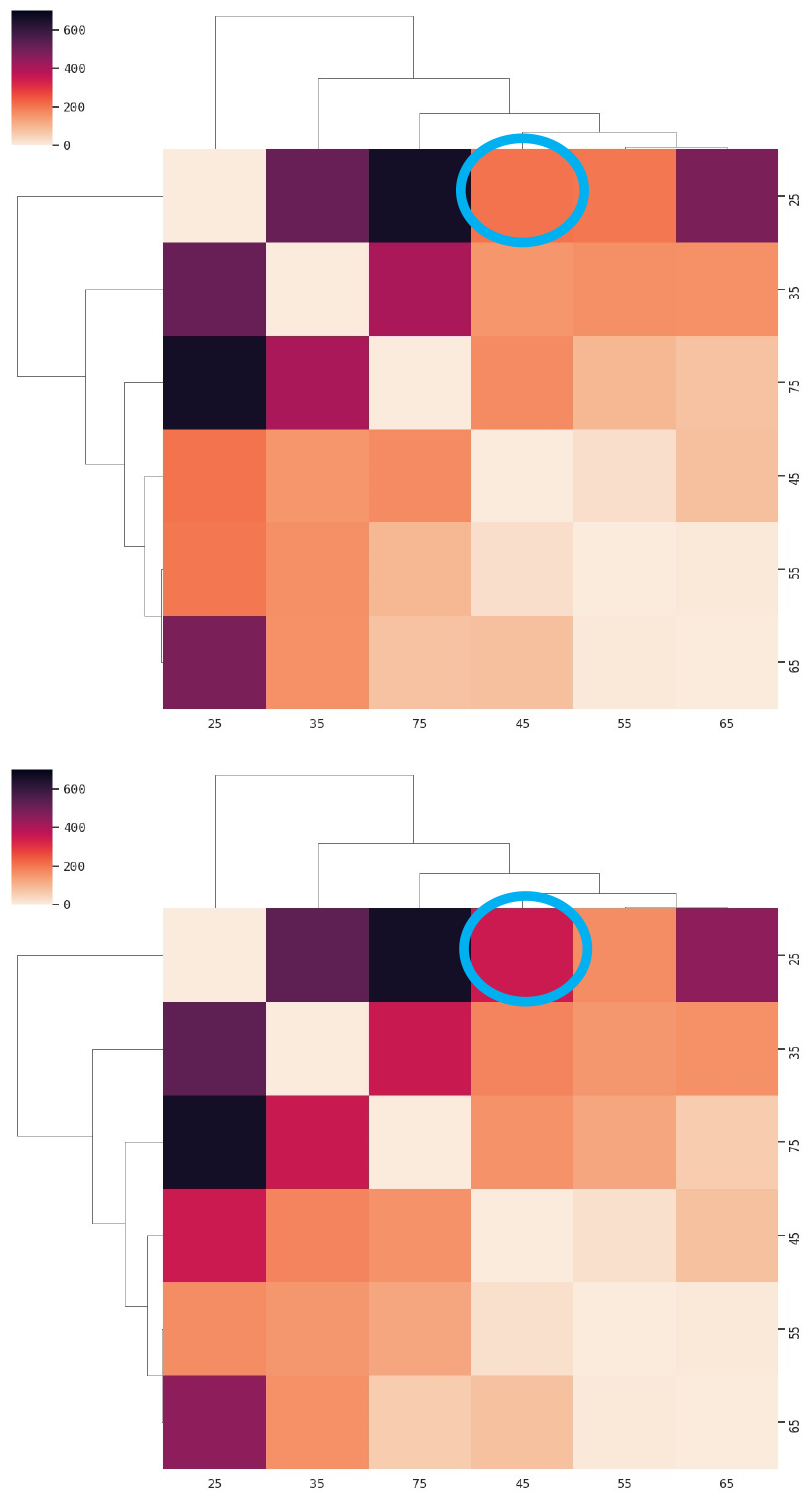
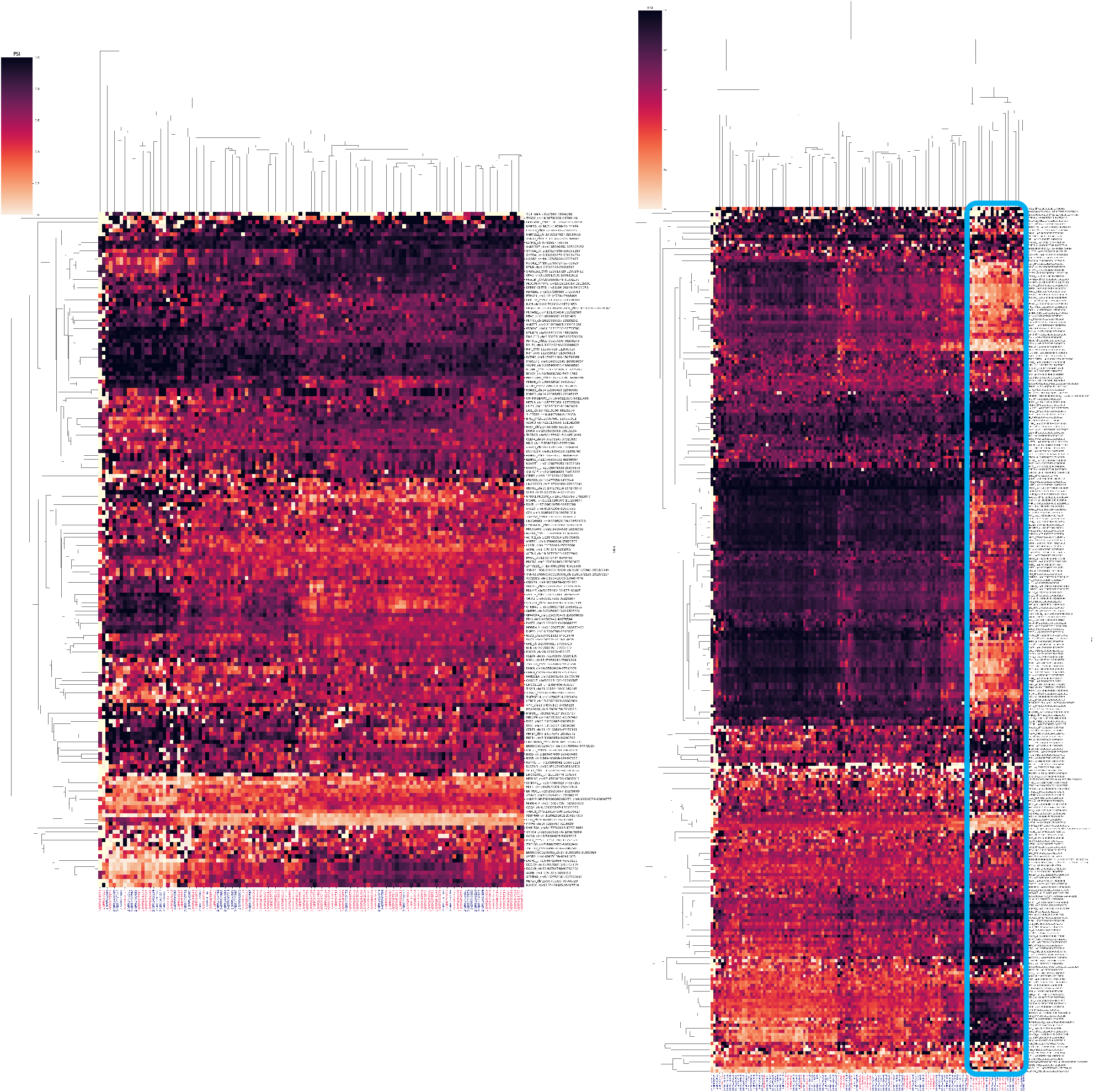
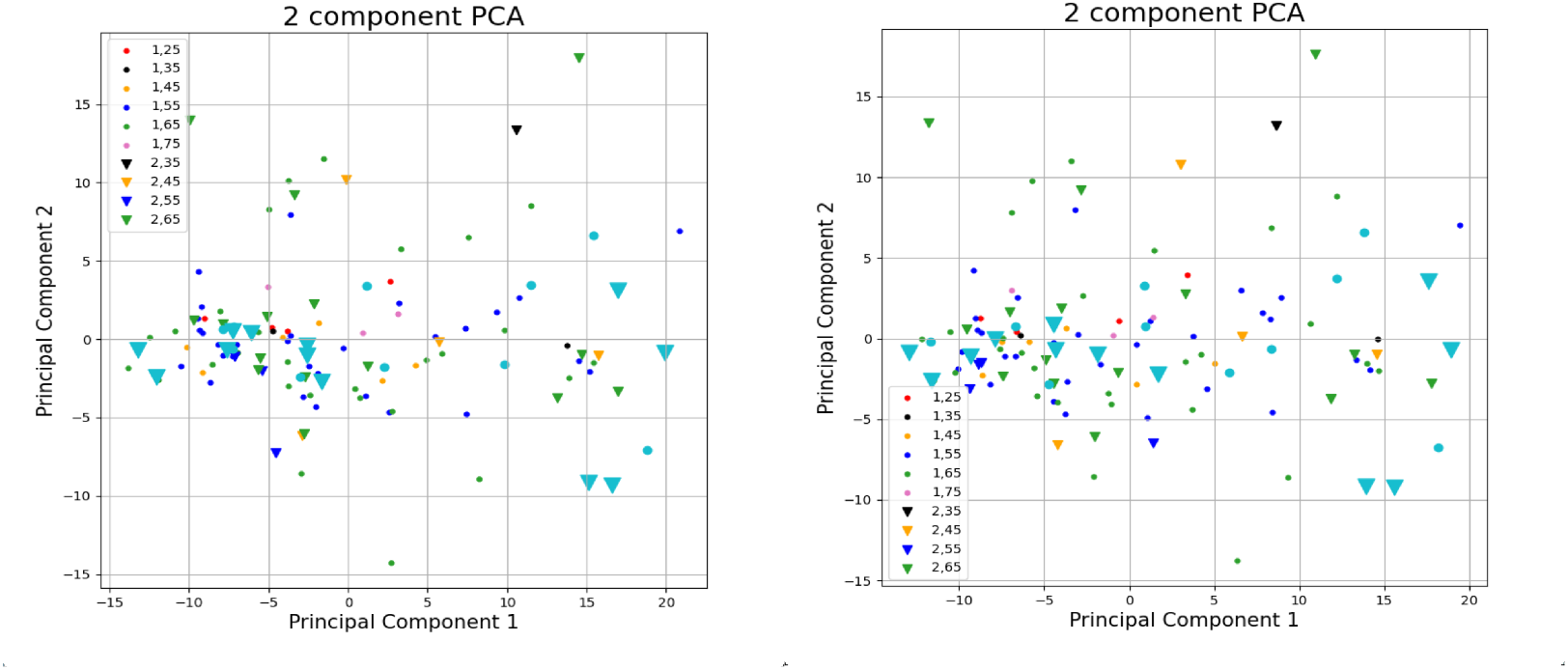
Differential splicing analyses of GTEx brain RNA-seq data with MntJULiP and Jutils. (A) Distance matrix of age group DSR comparisons, without and with ‘biological sex’ as covariate. (B) Jutils heatmaps of DSR events from the comparison between the female (‘F’) and male (‘M’) sample groups, without and with ‘age at death’ as covariate. PSI intron values estimated with MntJULiP and uploaded into Jutils were plotted, with rows (events) and columns (samples) clustered using the ‘weighted’ method and ‘cityblock’ similarity metric. The covariate-corrected heatmap shows a distinct cluster, marked with a box. (C) Jutils PCA plots from the comparison between female (‘F’) and male (‘M’) sample groups, without (left) and with (right) ‘age at death’ as covariate. PSI values estimated with MntJULiP and uploaded into Jutils were plotted using shapes to distinguish among condition groups (‘F’, ‘M’), and colors to mark subsets of samples with different covariate and attribute values. The subgroup of samples with a distinct splicing program is shown in cyan.

## Conclusions

Confounders such as sex, age and other biomedical attributes inherent to RNA-seq datasets from disease and population studies can bias bioinformatics analyses, yet tools that can effectively account for their effects are lacking. We accounted for covariate effects in differential splicing models in our tool MntJULiP and implemented supporting PCA visualizations in the Jutils suite. We show that they accurately and effectively remove covariate effects from both simulated and real data to uncover the true patterns of splicing variation. In particular, analyses of GTEx RNA-seq data from frontal cortex identify a distinct splicing program in a subgroup of individuals. Therefore, MntJULiP and Jutils are highly effective and efficient analysis tools for large-scale complex RNA-seq datasets with confounding factors, which can reveal new insights into disease and population.

## Acknowledgments

Computations were performed on the Advanced Research Computing at Hopkins (ARCH) computing facility supported by the National Science Foundation [OAC 1920103]. This work was supported in part by the National Institutes of Health [GM129085 to L.F].

*The authors declare that they have no conflict of interest*.

## Notes

https://github.com/splicebox/

### Competing Interest Statement

The authors have declared no competing interest.

