## Supplementary Figures and Tables for "MntJULiP and Jutils: Differential splicing analysis of RNA-seq data with covariates"

SUPPLEMENTARY MATERIAL FOR THE ARTICLE:
"MNTJULIP AND JUTILS: DIFFERENTIAL SPLICING ANALYSIS OF RNA-SEQ DATA WITH COVARIATES"
BY W.W. LUI, G. YANG and L. FLOREA

**Table of Contents:**


**Supplementary Methods**

**Methods M1.** Covariate augmented Bayesian models
**Methods M2.** Validation of covariate models on simulated data

**Supplementary Tables**

**Table S1.** Outline of simulation model for 2-way DSA and DSR comparisons.
**Table S2.** Outline of simulation model for multi(k=3)-way DSA and DSR comparisons.

**Supplementary Figures**

**Figure S1.** The MntJULiP intron-based feature models.
**Figure S2.** Evaluation of MntJULiP DSR function with covariates for pairwise comparison.
**Figure S3.** Evaluation of MntJULiP DSA function with covariates for pairwise comparison.
**Figure S4.** Evaluation of MntJULiP DSR (A-C) and DSA (D-F) functions with covariates for multi(k=3)-way comparison.
**Figure S5.** Alternative splicing profiles of brain tissues, from analysis of 1398 GTEx brain RNA-seq samples.
**Figure S6.** Jutils heatmaps of DSA events differentially spliced between the 'F' and 'M' categories.

### Supplementary Methods

#### Methods M1. Covariate augmented Bayesian models

*The differential splicing abundance (DSA) model*

This model tests an intron for differences in abundance among K conditions. Let N be the number of samples, K conditions and P covariates (including the comparison). We assume the read count y of intron v in sample i follows a negative binomial distribution NB($\mu_k$ + $\mathbf{x_i}\boldsymbol{\beta}_k$+ $a_k$, θ) with mean μ, K coefficient column vectors $\boldsymbol{\beta}_k$ of length P, sample intercept $a_k$, and N covariate row vectors $\mathbf{x_i}$ of length P. We consider a loose prior with an empirical $\hat{\mu}$ (the sample mean) modeled by a normal distribution: β ~ N(0, sqrt($\hat{\mu}$)) to effectively capture the variances across different conditions and within individual samples. Additionally, the dispersion parameter is restricted by a generic prior ϕ-1 ~ sqrt(N(0, 1)). Finally, to account for low expression genes and transcripts and low read counts, we introduce a zero inflated modifier. The zero inflated enhanced negative binomial (ZINB) Bayesian model is:

$$Y = f(x) = \begin{cases} 0, & \text{with probability } \pi \\ NB(\mu + x_i\beta_k + a_k), & \text{with probability } (1-\pi) \end{cases}$$

Maximum likelihood estimation is performed with respect to parameters μ, β, a, and θ, separately for the null and alternative models (*i.e.*, samples generated from a single-condition and from a K-condition experiment, respectively) to obtain log likelihoods $L(\theta_0)$ and $L(\theta_1)$ for testing [2].

*The differential splicing ratio (DSR) model*

The model tests introns within a 'bunch' for differences in splicing ratios among conditions. We assume the read counts $y_{i1}$, $y_{i2}$, …, $y_{iM}$ in sample i for a 'bunch' with M introns follow a Dirichlet multinomial distribution with concentration parameters $\alpha_{i1}$, $\alpha_{i2}$, …, $\alpha_{iM}$, the M coefficient row vectors $\boldsymbol{\beta}_m$ of length P, the intercepts $a_m$, N covariate column vectors $\mathbf{x}_i$ of length P, and 'bunch' total $\mathbf{n}_{iB} = \Sigma_{m'} y_{im'}$ [1]:

$$y_{i1}, y_{i2}, \ldots y_{iM} \,|\, \mathbf{n}_i \sim DM(\mathbf{n}_{iB}, \alpha_{i1}p_{i1}, \alpha_{i2}p_{i2} \ldots, \alpha_{iM}p_{iM})$$

$$p_{im} = \exp(\mathbf{x_i}\beta_m + a_m) \,/\, \Sigma_{m'} \exp(\mathbf{x_i}\beta_{m'} + a_{m'})$$

Maximum likelihood estimation is performed with respect to parameters α, β, and a, separately for the null and alternative models (excluding and including the condition column *x,* respectively) to obtain the log likelihoods $L(\theta_0)$ and $L(\theta_1)$ for testing [2].

*Covariate adjusted PSI and abundance estimates*

Using the covariate augmented models above, we generate new adjusted counts and PSI values per sample. After regressing out confounders, we obtain residuals *r,* applying a weakly informative prior r ~ N(0, 10). For *abundance* estimates, r is estimated for the alternative model with the optimized μ, β, a, and θ:

$$y \sim \begin{cases} 0, & \text{with probability } \pi \\ NB(\mu + x_i\beta_k + a_k + r, \theta), & \text{with probability } (1-\pi) \end{cases}$$

For splicing ratios (PSI) estimates, r is estimated for the alternative model using the optimized α, β, and a [1]:

$$y_{i1}, y_{i2}, \ldots y_{iM} | \mathbf{n}_i \sim DM(\mathbf{n}_{iB,} \alpha_{i1}p_{i1}, \alpha_{i2}p_{i2} \ldots, \alpha_{iM}p_{iM}).$$

$$p_{im} = \exp(\mathbf{x}_i\beta_m + a_m + r_i) / \Sigma_{k'} \exp(\mathbf{x}_i\beta_{m'} + a_{m'} + r_i)$$

The negative binomial and the Dirichlet multinomial models are implemented in PyStan, the Python package for Bayesian inference. Estimated counts were added to the output 'intron_data.txt' file, complementing the existing raw count values, and PSI values derived from the raw read counts and model-estimated PSI values were reported in a new file, 'group_data.txt'.

**Methods M2. Validation of covariate models on simulated data**

To validate the covariate models, we simulated data for one 'condition', with values 'disease' and 'control', with one covariate, 'biological sex', with values 'M' and 'F'. Changes in splicing were simulated as described in [1]. *For DSR*, we simulated 5 samples in each of the four categories 'condition'*x*'biological sex'. Starting from an empirical transcript expression matrix of lung fibroblast cells trained on the SRR493366 data set and GENCODE v.41 as reference, we generated 11.5 million 100 bp long paired-end reads per sample from 2000 genes with two or more expressed isoforms, using Polyester [2]. For changes due to the 'condition', 500 genes had the expression levels of the top two transcript isoforms swapped to simulate differential splicing (DS); for changes due to 'biological sex', a non-overlapping set of 200 genes were perturbed in the same way (see **Supplementary Table S1**). *For DSA*, differences due to 'condition' were simulated at 600 genes, with 200 differentially spliced only (DS), by swapping the isoforms' expression levels; 200 differentially expressed only (DE), by doubling or halving the expression of the gene and therefore of all its isoforms; and both differentially expressed and differentially spliced (DE-DS). Differences in 'biological sex' were marked as changes in 300 genes, 100 each from the DS, DE and DE-DS categories (see **Supplementary Table S2**).

**Supplementary Table S1.** Outline of simulation model for 2-way DSA (A) and DSR (B) comparisons.

(A)

| Condition\Covariate* | M | F |
|---|---|---|
| Control | 0 | 200x (DS) |
| Disease | 500d (DS) | 200x (DS) + 500d (DS) |

*Genes modified versus reference (M,Control): 500 genes differentiate based on disease status (‘d’), and 200 non-overlapping genes based on biological sex (‘x’).

(B)

| Condition\Covariate* | M | F |
|---|---|---|
| Control | 0 | 100x(DS)+100x(DE)+100x(DS+DS) |
| Disease | 200d(DS)+200d(DE)+200d(DE+DS) | 100x(DS)+100x(DE)+100x(DE+DS)<br>200d(DS)+200d(DS)+200d (DE+DS) |

*Genes modified versus reference (M,Control): 600 genes differentiate based on disease status (‘d’), and 300 non-overlapping genes based on biological sex (‘x’).

**Supplementary Table S2.** Outline of simulation model for mult(k=3)-way DSA and DSR comparisons. Genes marked as (DS) and (DE+DS) are used to evaluate the DSR function, and all categories (DS), (DE) and (DE+DS) are used in assessing the DSA function.

| Condition\Covariate* | M | F |
|---|---|---|
| Control | 0 | 100x(DS)+100x(DE)+100x(DE+DS) |
| Disease | 100d(DS)+100d(DE)+100d(DE+DS) | 100x(DS)+100x(DE)+100x(DE+DS)<br>100d(DS)+100d(DE)+100d(DE+DS) |
| Stage2 & Disease | 100ds2(DS)+100ds2(DE)+100ds2(DE+DS) | 100x(DS)+100x(DE)+100x(DE+DS)<br>100ds2(DS)+100ds2(DE)+100ds2(DE+DS) |
| Stage2 | 200s2(DS)+200s2(DE)+200s2(DE+DS) | 100x(DS)+100x(DE)+100x(DE+DS)<br>200s2(DS)+200s2(DE)+200s2(DE+DS) |

*Genes modified versus reference (M,Control): 1,200 genes differentiate based on disease status (‘d’- perturbed in ‘disease’ state only, ‘ds2’ – perturbed in both the ‘disease’ and ‘stage2’ states, and ‘s2’ – perturbed in ‘stage2’ only); and 300 non-overlapping genes based on biological sex (‘x’).

**Supplementary Figure S1.** The MntJULiP intron-based feature models. *Left*, differential splicing abundance (DSA): Each intron is analyzed individually, and the expression (abundance) level is compared between conditions C1,C2 and C3. *Right*, differential splicing ratio (DSR): Introns that share a splice junction ('bunch') are collectively analyzed, and the PSI values for the introns are compared between conditions C1 and C2. Shown are: an individual exon in a three-condition experiment, in the DSA diagram, and a three-intron 'bunch' in a two-condition experiment, in the DSR diagram.

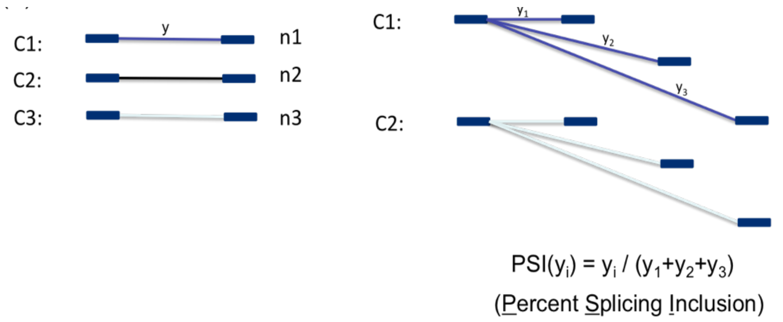


**Supplementary Figure S2.** Evaluation of MntJULiP DSR function with covariates for pairwise comparison, and by comparison to LeafCutter. (A) Performance evaluation: Sn=TP/(TP+FN), Pr=TP/(TP+FP), F-val=2*Sn*Pr/(Sn+Pr). (B) Breakdown of FPs by covariate versus extrinsic factors. (C) PCA plots of samples based on estimated PSI values, before and after covariate treatment.

(A)

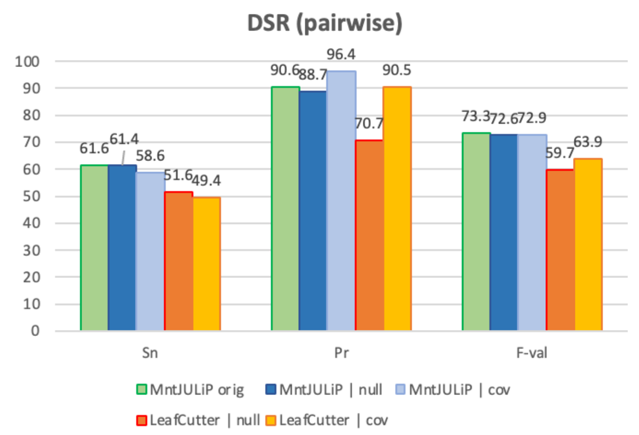


(B)

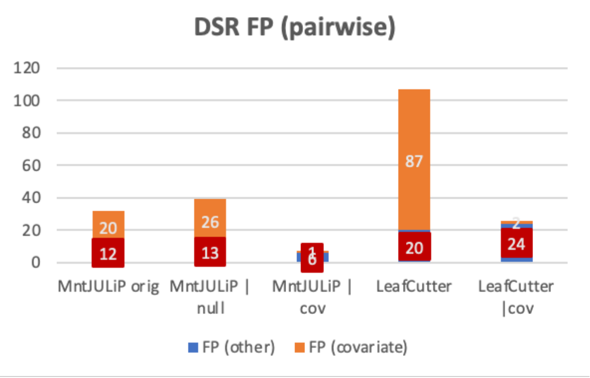


(C)

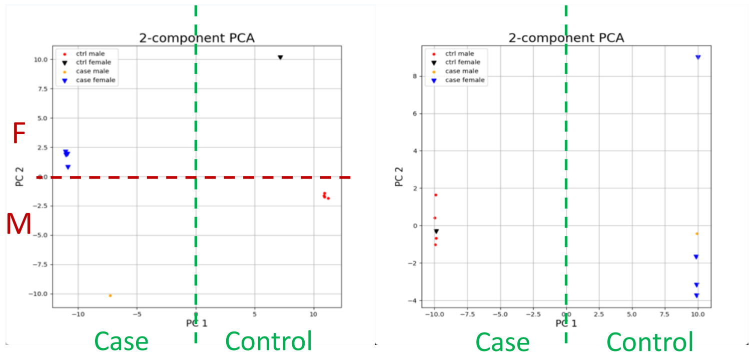

**Supplementary Figure S3.** Evaluation of MntJULiP DSA function with covariates for pairwise comparison. (A) Performance evaluation: Sn=TP/(TP+FP), Pr=TP/(TP+FN), F-val=2*Sn*Pr/(Sn+Pr). (B) Breakdown of FPs by covariate versus extrinsic factors. (C) PCA plots of samples based on estimated intron abundance, before and after covariate treatment.

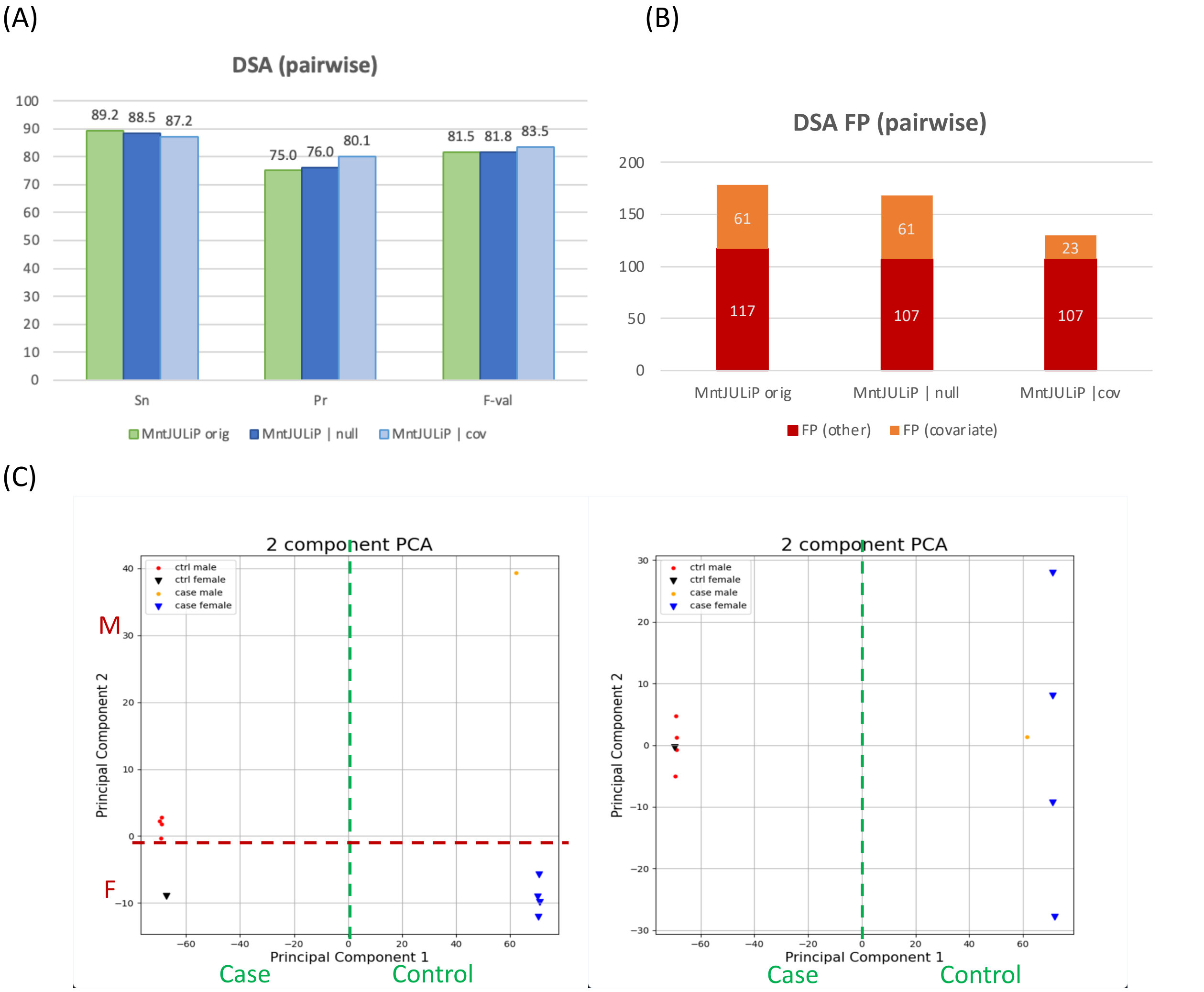

**Supplementary Figure S4.** Evaluation of MntJULiP DSR (A-C) and DSA (D-F) functions with covariates for multi(k=3)-way comparison. (A,D) Performance evaluation: Sn=TP/(TP+FN), Pr=TP/(TP+FP), F-val=2*Sn*Pr/(Sn+Pr). (B,E) Breakdown of FPs by covariate versus extrinsic factors. (C,F) PCA plots of samples based on estimated PSI values (C) and intron abundance (F), respectively, before (left) and after (right) adjustment for sex as covariate.

(A)

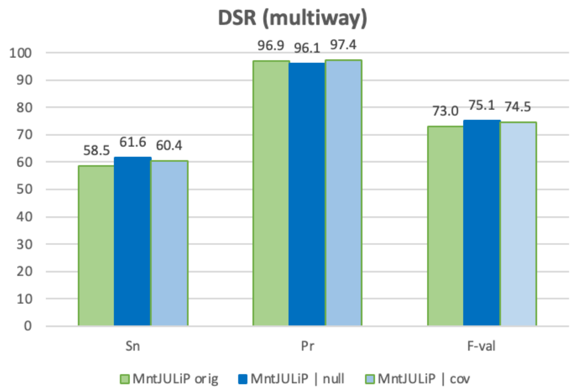


(B)

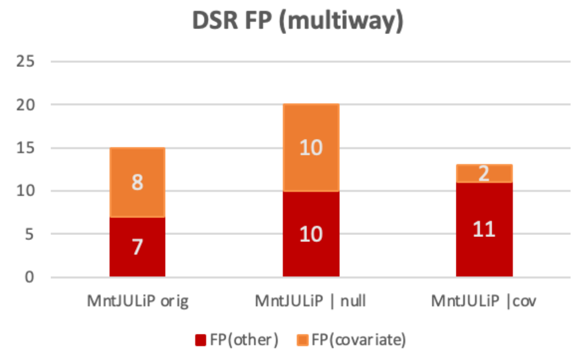


(C)

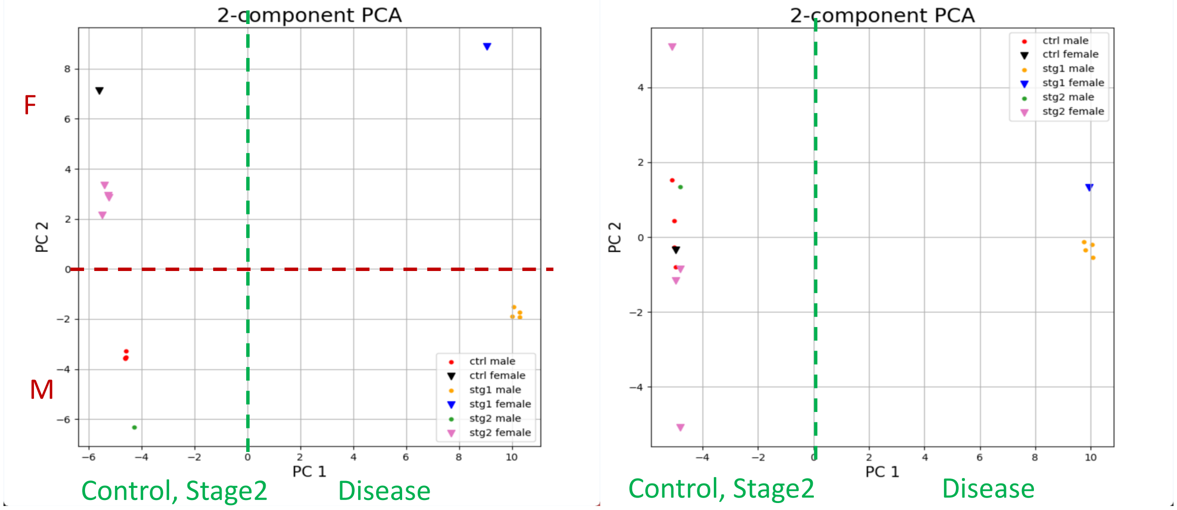


(D)

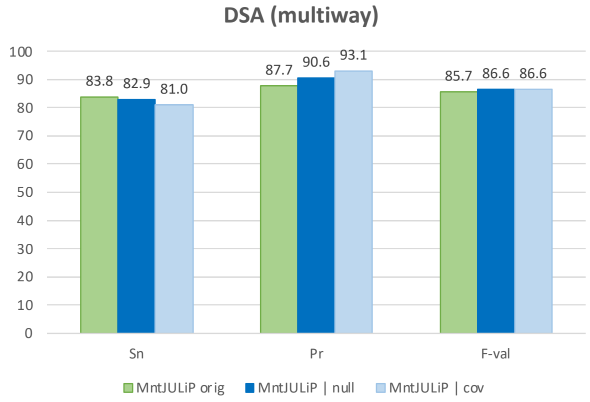


(E)

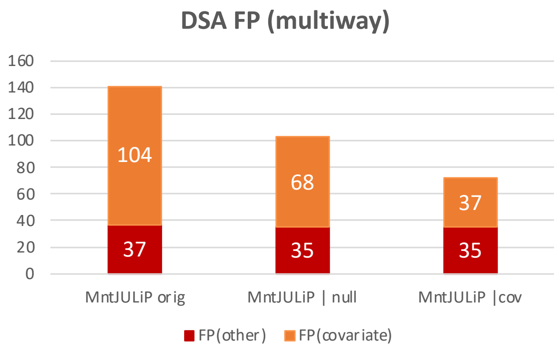

(F)

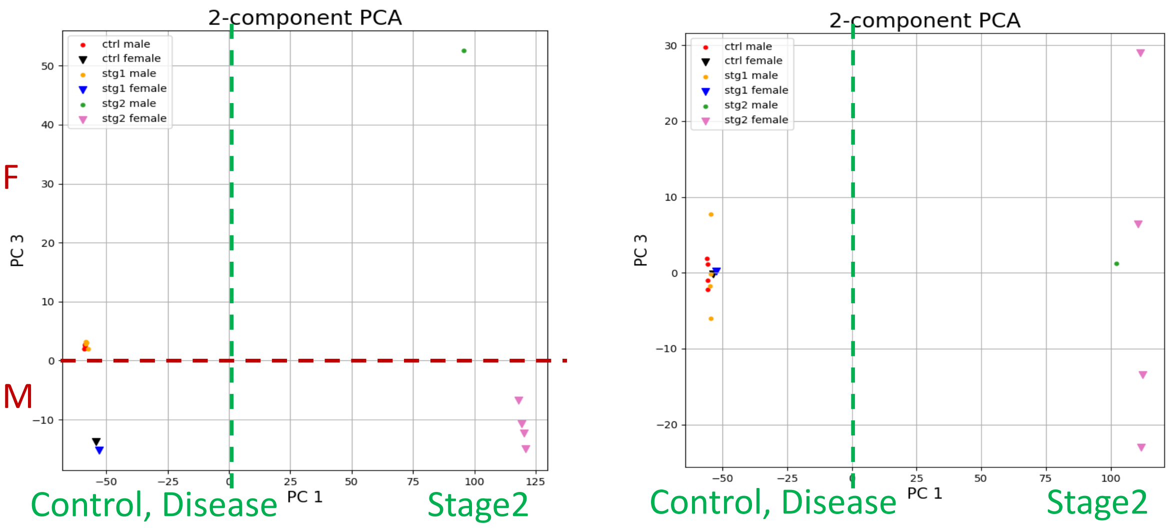


**Supplementary Figure S5.** Alternative splicing profiles of brain tissues, from differential splicing analysis of 1398 GTEx brain RNA-seq samples. (A) Distance matrices constructed from DSR differential splicing events: (left) without covariate treatment, (center) with 'biological sex' as covariate, and (right) with 'age at death' as covariate. (B) Similarly, for DSA differential splicing.

(A)

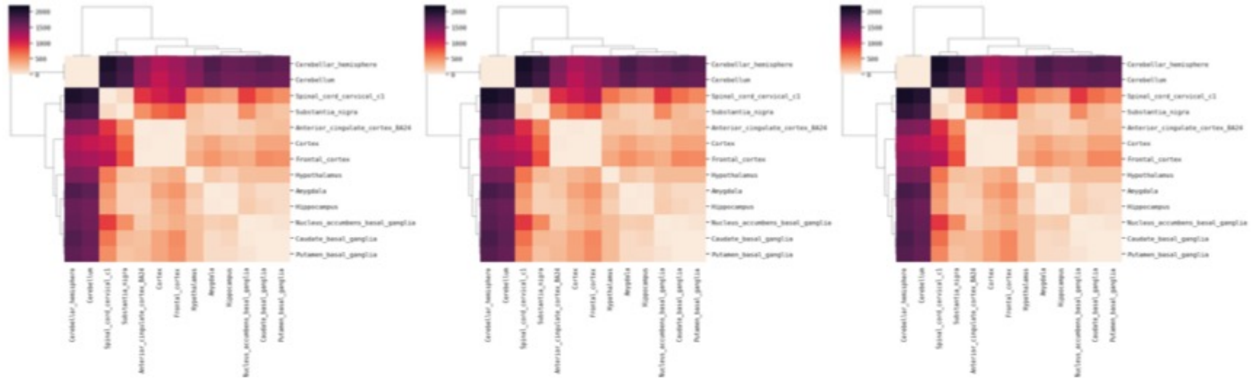

(B)

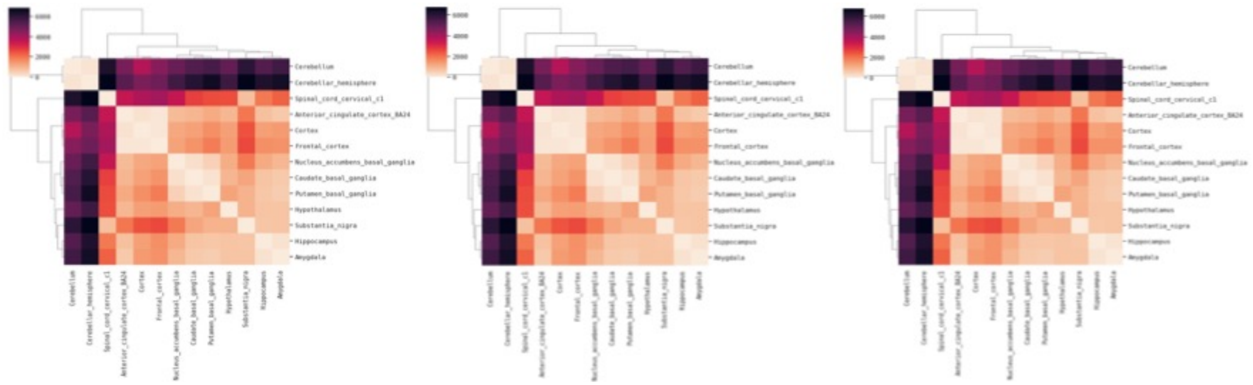

**Supplementary Figure S6.** Jutils heatmaps and PCA plots of DSA differentially spliced events (introns) between the 'F' and 'M' groups, without covariate treatment (left) and treating 'age at death' as covariate (right). In both scenarios, 'F' (marked with 1, dots) and 'M' (marked with 2, inverted triangles) form distinct clusters in both scenarios.

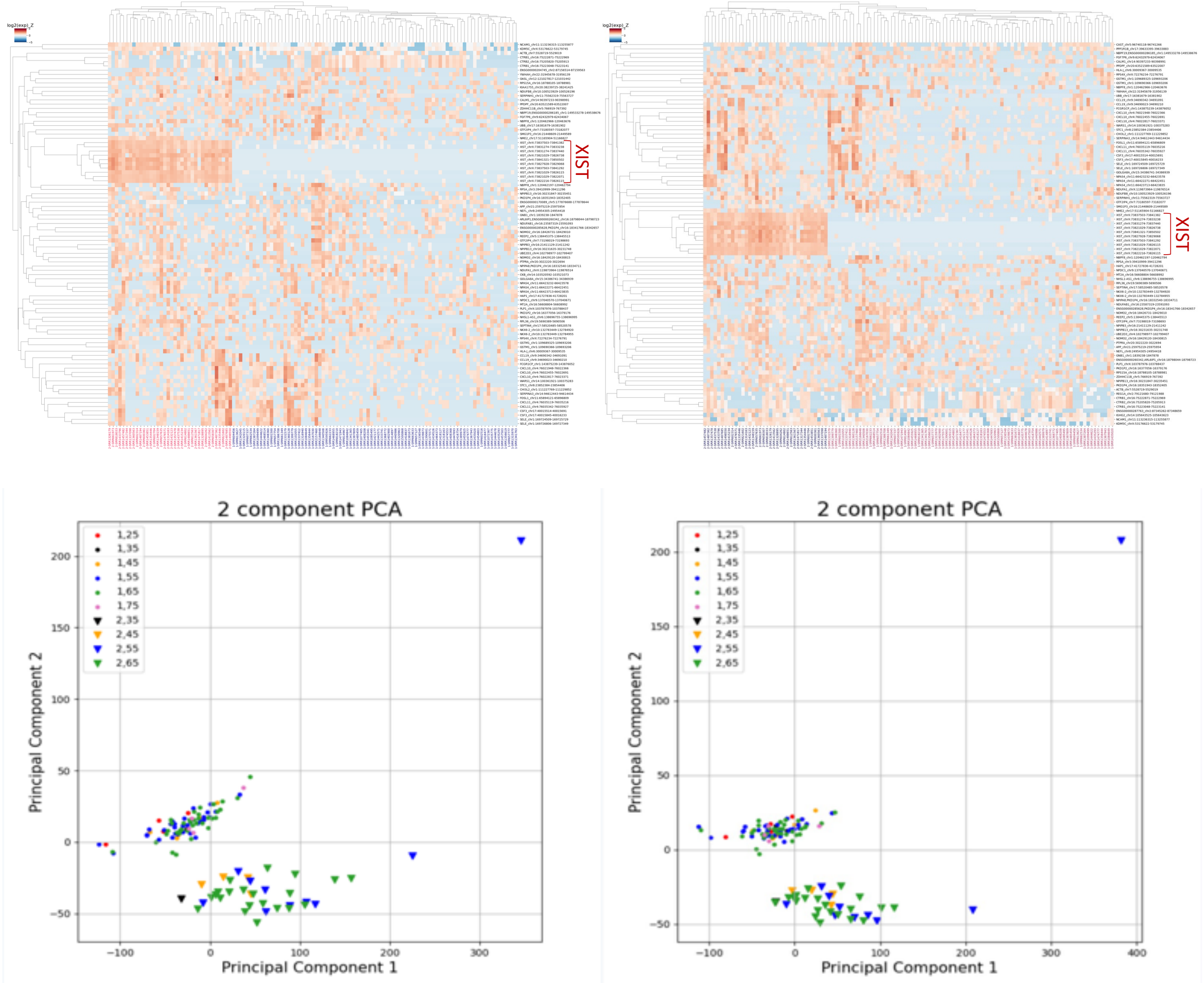
